# Body oscillations reduce the aerodynamic power requirement of wild silkmoth flight

**DOI:** 10.1101/2024.07.07.602433

**Authors:** Usama Bin Sikandar, Brett R. Aiello, Simon Sponberg

## Abstract

Insects show diverse flight kinematics and morphologies reflecting their evolutionary histories and ecological adaptations. Many silkmoths utilizing low wingbeat frequencies and large wings to fly display body oscillations: Their bodies pitch and bob periodically – synchronized with their wing flapping cycle. Similar oscillations in butterflies augment weight support and thrust and reduce flight power requirements. However, how the instantaneous body and wing kinematics interact for these beneficial aerodynamic and power consequences is not well understood. We hypothesized that the body oscillations affect aerodynamic power requirements by influencing the wing rotation relative to the airflow. Using three-dimensional forward flight video recordings of four silkmoth species and a quasi-steady blade-element aerodynamic method, we found that the body pitch angle and the wing sweep angle maintain a narrow range of phase differences to enhance the angle of attack variation between each half-stroke due to enhanced wing rotation relative to the airflow. This redirects the aerodynamic force to increase upward and forward force during downstroke and upstroke respectively thus lowering the overall drag without compromising weight support and forward thrust. A reduction in energy expenditure is beneficial because adult silkmoths do not feed and rely on limited energy budgets.

## 1 Introduction

The incredible variation of insect morphology and flight behaviors is a key factor in their wide diversification. One of the most ubiquitous flight behaviors is forward flight which insects use to explore, forage and migrate. To vary the forward speed, insects like hawkmoths, bees and flies primarily vary their body pitch angles to tilt their stroke planes. This positions the wingstroke-averaged aerodynamic force at a certain angle relative to gravity so that they can prescribe a forward speed while generating sufficient weight support [1]. Holding the bodies at a constant angle, they can turn their wings around the wing hinge (known as feathering or wing pitching) to precisely tune the angle of attack as well as redirect the aerodynamic force into usable directions over a wingstroke. However, in many butterflies and silkmoths, body pitch angle has been observed to continuously change during a wingstroke [2]–[4]. This makes the wings rotate relative to the incoming airflow and results in an oscillating stroke plane. Butterflies and silkmoths have also been observed to bob during each wingstroke [3]–[5].

Both pitching and bobbing oscillations complicate the control of effectively tuning the angle of attack and redirecting the aerodynamic force at the same time to propel forward and support body weight. These body oscillations, however, are a consequence of wing-body coupling dynamics that exist in all flapping insects but are particularly pronounced in butterflies with low wing loading and wingbeat frequency [6], [7]. However, it is not known how the magnitude of body oscillations is precisely associated with wingbeat frequency and wing loading or other parameters such as flapping amplitude. In addition, these parameters affect both aerodynamic and inertial forces. Thus it is not obvious which of these forces (or both) drives the body oscillations. Additionally, the energy expenditure in a flight style involving body oscillations seems higher because of the extra energy spent on the kinetic energy of oscillations. However, silkmoths have limited energy budgets because they do not feed as adults. Do they have a strategy to conserve power during flight? Other cases in biomechanics also exist where the active movement of one part of the body causes another part to passively oscillate to affect the energy requirements of locomotion. For example, in humans, passive arm swinging coupled with walking motion reduces metabolic energy requirements by 12% in comparison to volitional steady arm holding and by 26% in comparison to anti-phase arm swinging [8]. This inspires us to explore how the aerodynamic power requirements would be affected in the absence of body oscillations.

In this study on silkmoth flight, we aim to answer four questions to present a more holistic understanding of the aerodynamic implications of body oscillations. First, we investigate which wing movement and morphological parameters are associated with the magnitude of body oscillations in the kinematic modeling fits of four silkmoth species used in this study. Previous studies hint at the potential role of wingbeat frequency, flapping amplitude and wing loading but their precise relationship is not known. This analysis is meant to establish exemplars with different degrees of body oscillations and wing kinematics rather than specifically comparing the full flight kinematics of the four species. Second, we investigate whether inertial or aerodynamic forces are the main drivers of silkmoth body oscillations through wing-body coupling using a quasi-steady blade-element method. In birds, the generation of body oscillations is dominated by inertial forces from wing flapping [9]. However, butterfly studies show that body oscillation can be explained by the influence of the flapping wing aerodynamics on the body [6]. Third, we explore the aerodynamic consequences of body oscillations, given the pattern of the species-averaged wing and body kinematics observed. We analyze how observed body oscillations affect the wing angle of attack, angle of inclination and relative airflow speed relative to the horizontal. The angle of attack and relative airflow speed prescribe the aerodynamic force while the angle of inclination specifies the direction of the force. These quantities must be controlled for successful flight. Fourth, we explore the effect of coupling between body oscillations and flapping wing motion on flight performance in terms of weight support, forward thrust and aerodynamic power. We specifically ask whether there is a desirable phase difference between body oscillations and flapping wing motion, where they improve flight performance in terms of these parameters. Or the performance deteriorates compared with a hypothetical silkmoth body dynamics model that does not oscillate, irrespective of the phase difference. To answer these questions, we recorded 3-D forward flight kinematics of silkmoths flying in a wind tunnel. After extracting the useful kinematic measurements, we calculated the aerodynamic forces and power using our quasisteady blade-element method to analyze the dynamics and performance of their flight. Based on our kinematic measurements and aerodynamic calculations, we answer the four questions and then discuss their relevance to flight performance and behavior.

## 2 Methods

### 2.1 Body and wing measurements and morphometrics

In this study, we used live specimens of four wild silkmoth species that were available and occupy a large range of wingbeat frequency and wing size. The four species were *Automeris io* (1 specimen, 2 wingstrokes), *Actias luna* (3 specimens, 1 wingstroke each), *Antheraea polyphemus* (3 specimens, 1 wingstroke each), and *Hyalophora euryalus* (1 specimen, 4 wingstrokes). The body and wing morphologies were captured for every live specimen using our earlier techniques [4] through StereoMorph (v. 1.6.2) [10] within R software (v. 3.4.2). After each specimen had flown, its total body weight (*m*_*t*_) was recorded. The morphological data were subsequently processed in Matlab (v. R2018b - 9.5.0.944444) as per [4], [11]. To create a combined wing shape from the intersection of the fore- and hindwings, we adjusted the forewing’s long axis to make it perpendicular to the body’s long axis. The hindwing was maintained in its natural position, the state it is in when the wings spread wide and the moth is idle. This orientation was chosen after examining silkmoth flight videos where the hindwing’s long axis is consistently oriented posteriorly and parallel to the body’s long axis. All the wing morphology parameters utilized for aerodynamic analysis were derived from this combined wing, as described by [12].

### 2.2 Moth free-flight high-speed recordings

Live specimens were filmed while flying forward at a speed of 2-3 ms^−1^ in a wind tunnel. We carried out free-flight experiments within a 100 × 60.96 cm segment of an open-circuit Eiffel-style wind tunnel provided by ELD, Inc. from Lake City, MN, USA. For a detailed description of the tunnel, refer to [13]. Moths were prompted to fly at a wind velocity of 0.7 ms^−1^. We captured the flight sequences of silkmoths at 1000 frames s^−1^ using three coordinated high-speed cameras (resolution: 1280 × 1024 pixels), specifically the Mini UX 100 from Photron, San Diego, CA, USA. The wind tunnel’s lighting consisted of six 850 Nm infrared lamps from Larson Electronics, Kemp, TX, USA, alongside a neutral density-filtered LED ‘moon’ light (Neewer CW-126) [14]. The footage was then processed and standardized in XMALab [15]. We identified and marked seven critical points on the moth which include the head’s rostral tip (situated between the antennae), the thorax-abdomen intersection, the abdomen’s rear end, left and right forewing attachment points, the right wing tip, and the intersection point of fore- and hind-wings on the distal edge of the right wing. These seven points were clearly identifiable and thus could be consistently tracked across species.

### 2.3 The quasi-steady aerodynamic model

The time series of the seven tracked points in 3-D were used to extract 3-D body velocities (*u, v* and *w*), body pitch angle *χ* and wing kinematic angles using custom MATLAB code as previously done in [11]. The wing kinematic angles used to specify wing motion were sweep angle *ϕ*, deviation angle *θ*, feathering angle *α* and stroke-plane angle *β* using a convention similar to [16]. The extracted time series of each kinematic parameter over each wingstroke was fitted with a third-order Fourier series. By averaging the wing shapes and the Fourier-fitted time-dependent kinematics across all wingstrokes for each moth individual, we determined one representative set of wing shapes, sizes, and kinematics for each species. We then calculated the species-specific aerodynamic forces through our quasi-steady blade element model previously published [4]. This model calculates the cumulative aerodynamic force by considering the forces from translational and rotational movements of the wing [11], [17] as well as the force resulting from added mass [11], [18]. The translational forces were calculated based on the lift and drag coefficients of the hawkmoth (*M. sexta*) [19]. The equations of these forces acting on a small wing strip of width *dr* are as follows.

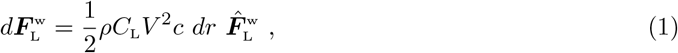

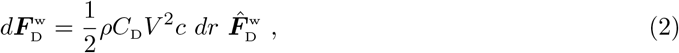

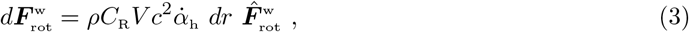

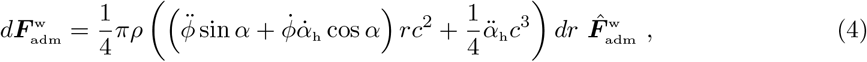

where *ρ* is the air density, the aerodynamic coefficients *C*_L_ and *C*_D_ were taken from [19], *V* is the relative airflow speed of the strip, *c* and *dr* respectively are chord length and width of the strip. Chord length *c* varies with *r* along the spanwise direction, the rotational aerodynamic coefficient 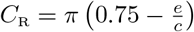 and *e* is the distance between the leading edge and wing pitching axis. The wing inclination angle, *α*_h_, is the angle the wing surface is inclined relative to the absolute horizontal plane, where *α*_h_ ≈ *α* − *β*, and its derivative represents the angular velocity of the wing pitching rotation in the global frame [1], [20]. Force on a small wing strip of length *dr* is integrated over the entire wing to calculate the total aerodynamic force on the wing.

A detailed formulation of our blade element model can be found in [4]. Such a first-principles model offers flexibility in directly examining the effects of various combinations of wing shapes and movements on aerodynamic force and power across a range of species. This approach lets us delve into how individual force components (like translational, rotational, and added-mass) shape the comprehensive aerodynamic force profile. Additionally, the force due to wing inertia was computed.

As measured in the body-attached frame, the inertial force on the body due to a wing strip of width *dr* flapping with angular velocity ***ω***_w_ at a distance *r* from the hinge is

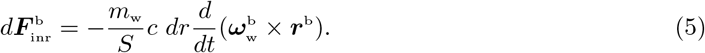

Because inertial forces are internal to the moth’s body, they do not generate a net force during steady wingstrokes. However, the aerodynamics could be indirectly influenced by the wing’s inertial forces (and the associated torques) through their effect on the body’s movement. That said, since wing kinematic measurements were taken relative to the stroke plane which is assumed fixed relative to the body, and all forces were converted to a frame attached to the body, any movement between the wings and body caused by inertial forces and its subsequent impact on aerodynamics are inherently accounted for. Next, the instantaneous aerodynamic power dissipated due to drag on a small wing strip of length *dr* is calculated as the work done by the drag force calculated using the equation

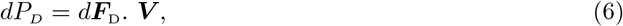

where *d****F***_D_ is the drag force on the wing strip and ***V*** is the airflow velocity incident on the wing strip.

### 2.4 Comparative models of body kinematic configuration

To determine the effects of body oscillations and their coupling with wing flapping on the aerodynamic performance of silkmoth flight, we use three model configurations of body kinematics to evaluate the aerodynamic forces and power, as shown in Fig. 1. Model 1 is the time-varying body kinematics, as they were measured. Model 2 is the wingstroke-averaged values of *χ, β, u* and *w*, representing hypothetical flight with no body oscillations. Model 3 is the same time-averaged body kinematics as Model 1 but time-shifted by a half time period, to introduce a 180^°^ phase shift relative to the wing kinematics. Model 3 also represents a hypothetical flight but the body kinematics in this case are anti-phase compared to the actual flight. The wing kinematics were the same time-varying periodic functions across these models. Differences between aerodynamics corresponding to Models 1 and 2 would inform us of the aerodynamic effects of having body oscillations versus not having body oscillations. Those between Models 1 and 3 would elicit whether the natural phase difference observed in steady forward flight has positive or negative effects on flight performance.

**Fig. 1.**
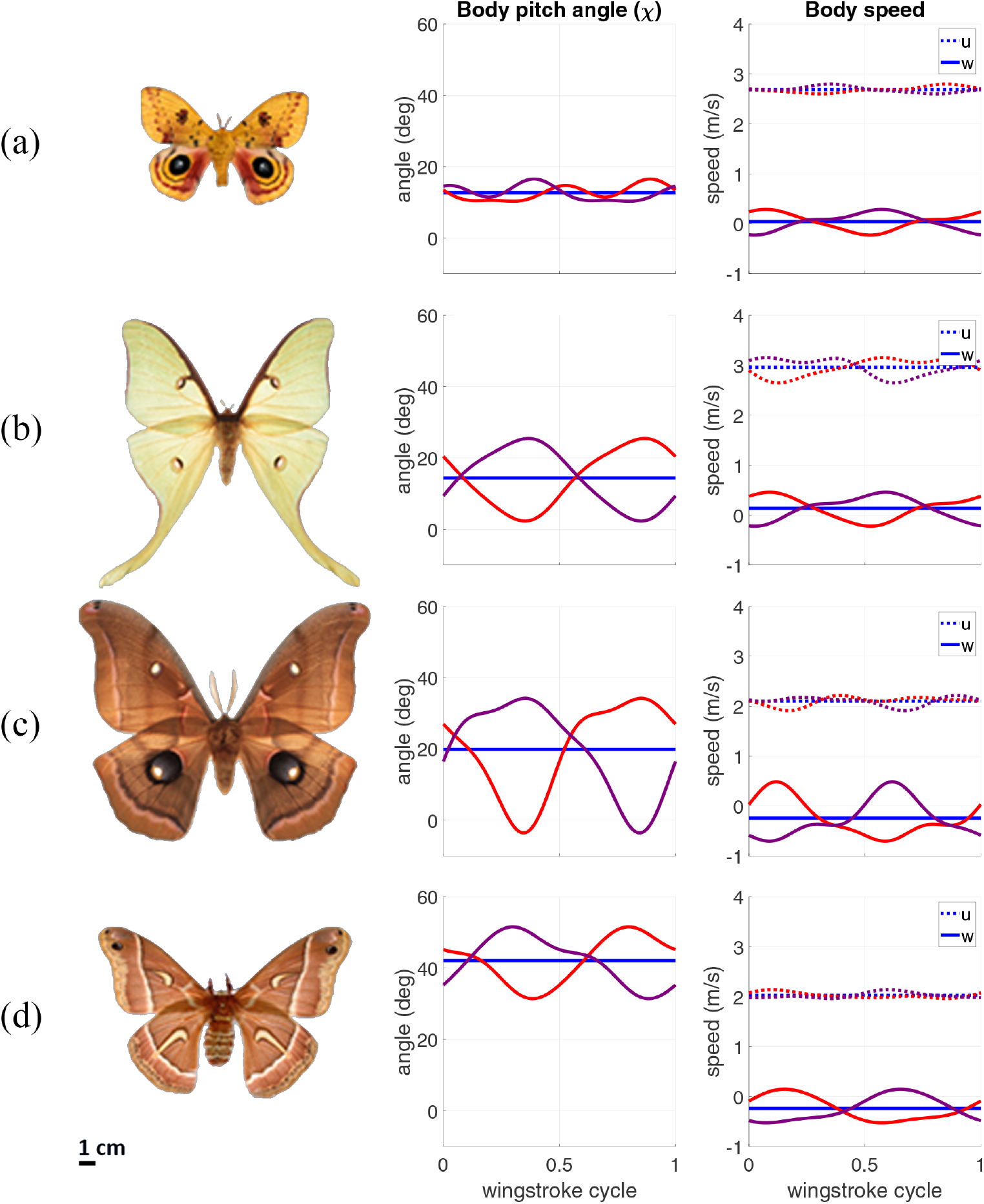
Panels a-d showing the three model configurations of body kinematics of four silkmoth species used in this study: (a) *A. io*, (b) *A. luna*, (c) *A. polyphemus* and (d) *H. euryalus*. The kinematics plotted are representative of each species and are calculated as the mean kinematics across all wingstrokes of a species. Model 1 (red) represents the time-varying configuration of body kinematics. Model 2 (blue) is the wingstroke-averaged version of body kinematics. Model 3 (purple) is Model 1 kinematics shifted by a half wingstroke period to introduce a 180^°^ phase shift relative to flapping wing motion.

## 3. Results

### 3.1 The amplitude of body oscillations varies with wingbeat frequency and wing loading

First, we identified the morphological and wing kinematic parameters underpinning variation in body oscillation magnitude. We plotted the amplitudes of body pitch angle *χ* and the body’s vertical velocity oscillations against several morphological and wing kinematic parameters. *A. io* displayed a noticeably smaller amplitude of body pitch oscillations compared to the other three species. The amplitude of body pitch *χ* oscillations of *A. io* was 6.9^°^. However, for the other three species, the amplitude of *χ* ranged between 21.8^°^ and 38.5^°^. The amplitude of oscillation in vertical velocity *w* was also the lowest for *A. io* at 0.5 ms^−1^, while the amplitude of *w* for other species ranged between 0.67 and 1.19 ms^−1^. Across the four silkmoth species, we found that the amplitude of body pitch angle *χ* oscillations correlate with wingbeat frequency (Fig. 2a) and the product of wingbeat frequency and wing loading (Fig. 2c). This means that silkmoths with lower wingbeat frequencies, smaller body mass and larger wings have larger oscillations in body pitch during forward flight. Extrapolating from this small dataset to all silkmoths will require a more extensive comparative study. Our goal here is to explore the aerodynamic consequences of variation in bobbing and pitching. The species sampled show substantial variation in these parameters justifying their use as exemplars for our aerodynamics study.

**Fig. 2.**
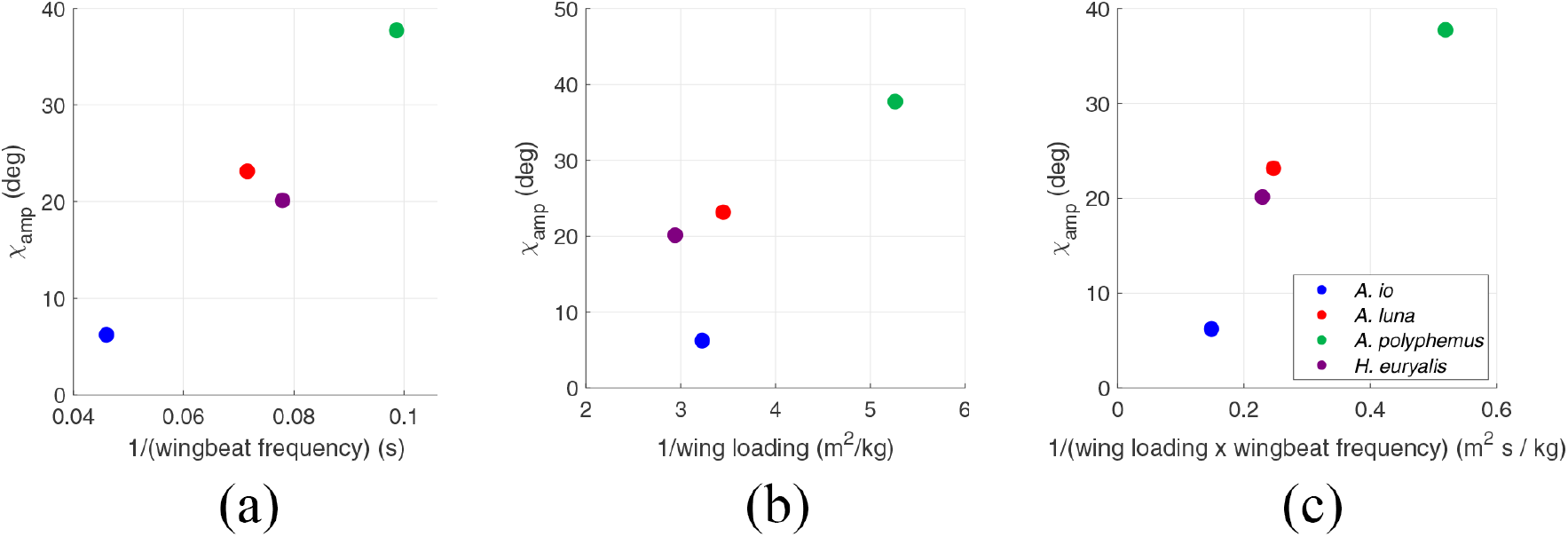
Scatter plots showing how the amplitude of body angle *χ* is associated with (a) wingbeat frequency, (b) wing loading and (c) the product of wingbeat frequency and wing loading. There is a correlation in (a) between the amplitude of *χ* and wingbeat time-period (1/wingbeat frequency): *r* = 0.97, *p <* 0.05. There is also a correlation in (b) between the amplitude of *χ* and wing loading (*W*_*s*_), *r* = −0.79, *p >* 0.05. Overall, there exists a correlation between the amplitude of *χ* and the inverse of the product of wing loading and wingbeat frequency: *r* = 0.94, *p <* 0.05.

### 3.2 Both aerodynamic and inertial forces contribute to body oscillations in silkmoths

Using these four species as exemplars, we next explore the source of body kinematic oscillations using quasi-steady blade-element method to determine whether aerodynamic or inertial force dominates in body oscillations. An important advantage of first principles physics models of flight is our ability to separate out specific force components. To do this, we compared the normalized aerodynamic and inertial forces (Fig. 3) generated by the observed species-averaged flapping wing motion and body kinematics in Fig. 1 (Model 1). Comparing the shapes and magnitudes of the two waveforms, we find that aerodynamic and inertial forces have comparable magnitudes, and thus, both forces contribute to silkmoth body oscillations. Therefore, contrary to in birds where inertial forces dominate, and in butterflies where aerodynamic forces drive body oscillations [6], [9], we found that both aerodynamic and inertial forces contribute to body oscillations in silkmoths.

**Fig. 3.**
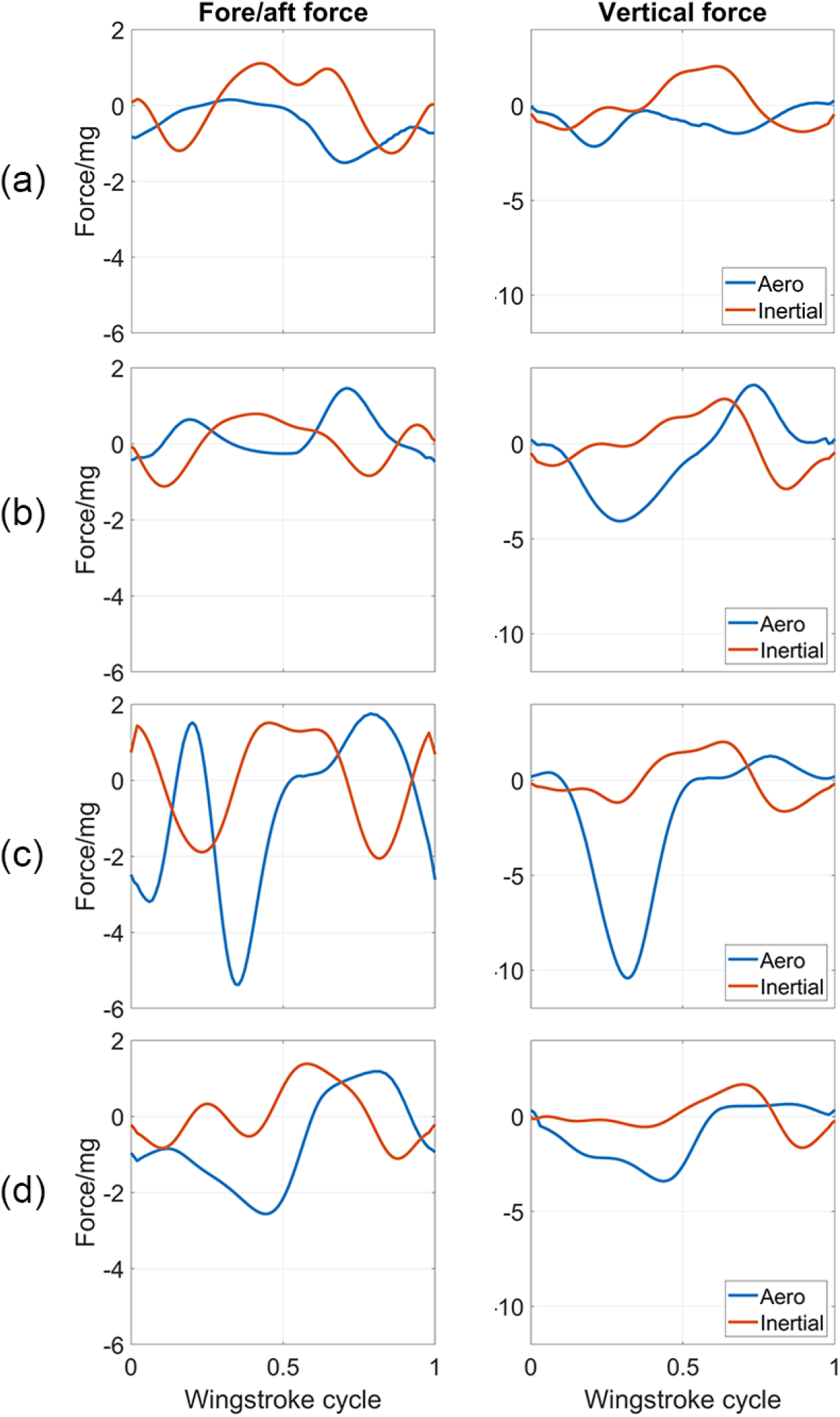
Panels a-d showing the normalized aerodynamic and inertial forces generated due to wing movement estimated for the four silkmoth species: (a) *A. io*, (b) *A. luna*, (c) *A. polyphemus* and (d) *H. euryalus*. The normalization makes the forces physically equivalent to nondimensional acceleration as multiples of g. Both aerodynamic and inertial forces have comparable magntiudes but aerodynamic forces dominate in some cases, such as, for *A. polyphemus*.

### 3.3 Body oscillations enhance weight support and forward thrust

Next, we determined the effects of body oscillations on the aerodynamic performance of flight. First, to understand why body oscillations affect aerodynamics, we plotted wing inclination angle *α*_h_, angle of attack *α*_v_ and relative airflow speed *v* for kinematics Models 1, 2 and 3 of all four silkmoth species (Fig. 4). From our blade element model, we know that these three parameters strongly modulate the aerodynamic force vector which is roughly perpendicular to the wing surface. The aerodynamic force must provide enough weight support and forward thrust for successful forward flight. Thus, the force vector can be resolved into two components: positive/negative weight support and forward/backward thrust. The angle of attack *α*_v_ and the relative airflow speed *v* specify the magnitude of the force while the wing inclination angle *α*_h_ specifies the relative size of weight support and thrust components.

**Fig. 4.**
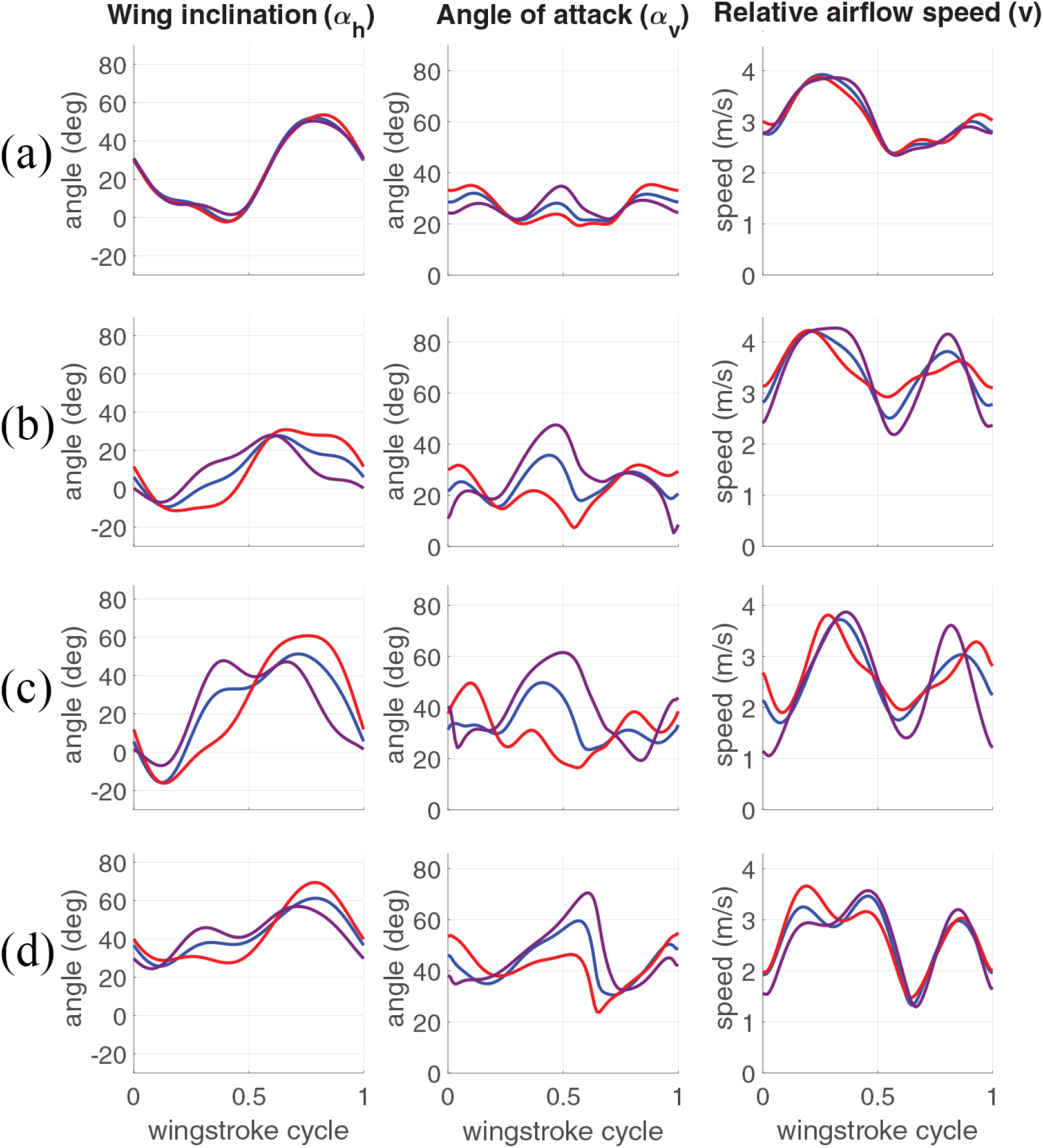
Graphs showing measured waveforms of wing inclination angle *α*_h_, wing’s mean (strip-averaged) angle of attack *α*_v_ and relative airflow speed *v* for each species: (a) *A. io*, (b) *A. luna*, (c) *A. polyphemus* and (d) *H. euryalus*. Model 1 (red) represents the time-varying configuration of body kinematics. Model 2 (blue) is the wingstroke-averaged version of body kinematics. Model 3 (purple) is Model 1 kinematics shifted by a half time-period to introduce a 180^°^ phase shift relative to flapping wing motion.

Comparing Models 1 and 2 (Fig. 4), we found that body oscillations, particularly in pitch angle *χ* modulate the time variation in *α*_h_. This results in, on average, a low *α*_h_ during the downstroke and a high *α*_h_ during the upstroke as shown in *α*_h_ panels of Fig. 4(a)-(d). A low *α*_h_ during the downstroke makes the aerodynamic force vector more vertically upward. This means that most of the aerodynamic force provides weight support rather than backward thrust, which drives forward flight. During the upstroke, a larger *α*_h_ directs the aerodynamic force more horizontally forward. This means a larger portion of the aerodynamic force provides a forward thrust rather than a downward force. Thus, given the species-averaged kinematics that we measured, enhanced weight support and forward thrust over the first and second half strokes respectively reduce the magnitude of the aerodynamic force required for flight.

The oscillations in *χ* not only modulate the relative sizes of weight support and thrust through appropriate variations in *α*_h_, they also affect the wing angle relative to the incoming airflow, thus modulating the wing angle of attack *α*_v_. As shown in panels *α*_v_ of Fig. 4(a)-(d), *α*_v_ in Model 1 stays lower than Model 2 for most of a wingstroke duration thus decreasing the drag force. But it does not go too low to lose the required lift force. Compared to the case of no-oscillations in Model 2, this results in a smaller overall force required for flight, thus reducing the power requirements.

### 3.4 Phase-coupling between body oscillations and wing flapping reduce the aerodynamic power requirements

To explore the existence of phase coupling between body oscillations and flapping wing motion, possibly as a preferred phase difference between the two, we calculated the phase difference between *χ* and wing sweep angle *ϕ* across all 10 wingstrokes from the three silkmoth species with large *χ* oscillations and scattered the values on a polar plot shown in Fig. 5. We found that the phase difference stays within a narrow range (30^°^-120^°^) spanning only about 90^°^, hinting at a possible phase coupling between body oscillations and flapping wing motion in silkmoth species whose body oscillations affect their flight aerodynamics. These values of phase difference during steady forward flight may be favorable in reducing aerodynamic power requirements.

**Fig. 5.**
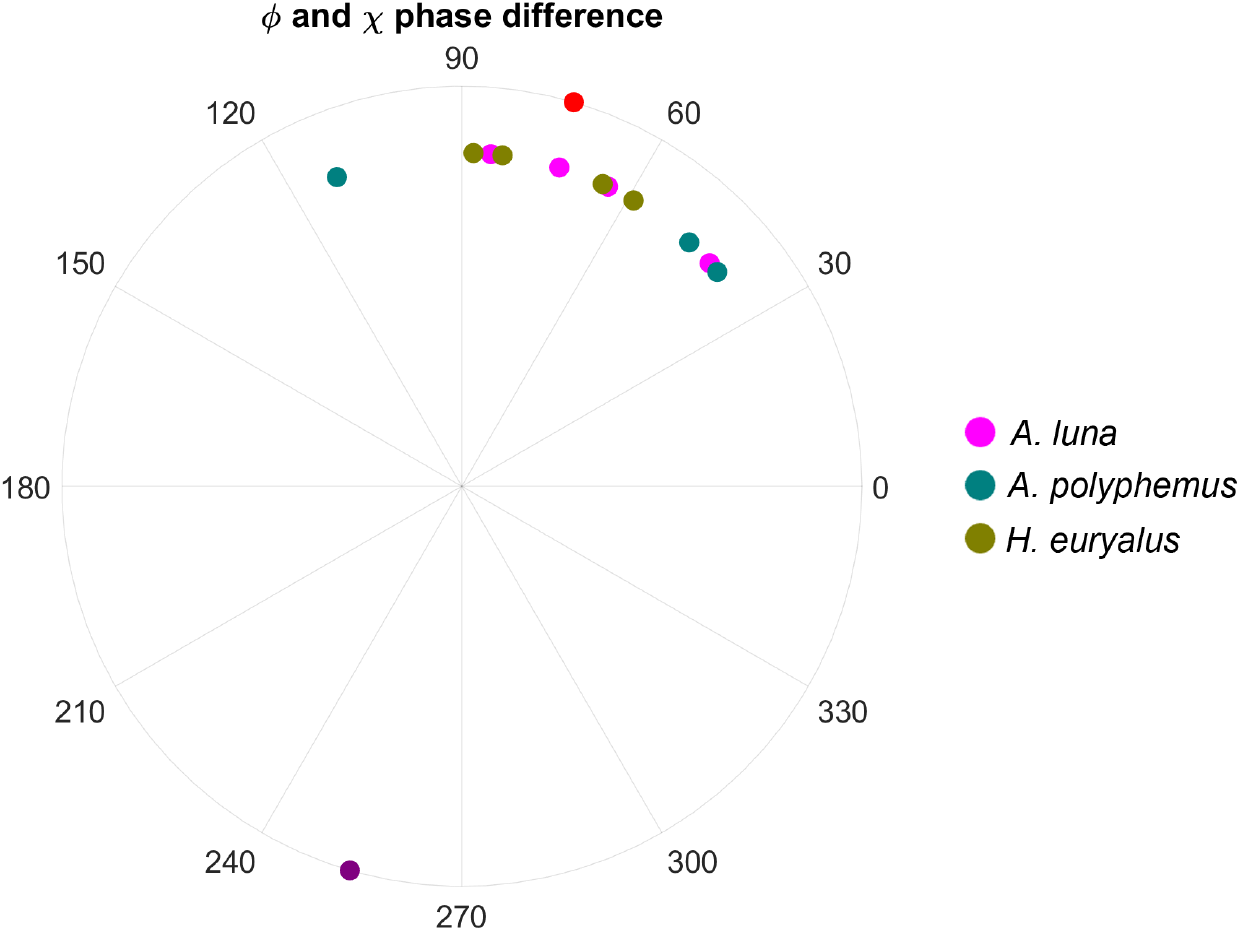
Scatter plot shows phase differences between body pitch angle *χ* and wing sweep angle *ϕ*. Red and purple markers on the circumference show phase differences of Models 1 and 3 respectively of the averaged representative wingstroke of *H. euryalus*. Each of the other markers shows a single measured wingstroke of silkmoths *A. luna, A. polyphemus* and *H. euryalus*.

To explore the aerodynamic implications of phase coupling between body oscillations and wing flapping, we compared the aerodynamic performance corresponding to Models 1 and 3, where Model 3 represents the kinematics in which body oscillations are phase-shifted by 180 degrees relative to flapping wing motion. Since silkmoth *A. io* did not show a considerable magnitude of body oscillations, we found that except for this species, both mean and peak aerodynamic power required to overcome drag increased in Model 3 compared to Model 1. As shown in Fig. 6, the increase in this mean power requirement ranges from 1.24 times for *H. euryalus* to 2.20 times for *A. polyphemus*, and the increase in peak power ranges between 2.15 times for *H. euryalus* and 3.74 times for *A. polyphemus*. Another important aerodynamic performance metric, lift-to-drag ratio shown in Fig. 6, is a measure of aerodynamic efficiency and it is calculated as the ratio between the wingstroke-averaged lift and drag coefficients. In silkmoths *A. luna* and *A. polyphemus*, the lift-to-drag ratio is notably higher for Model 1. This further emphasizes the role of body oscillations and their coupling with wing motion in determining aerodynamic efficiency.

**Fig. 6.**
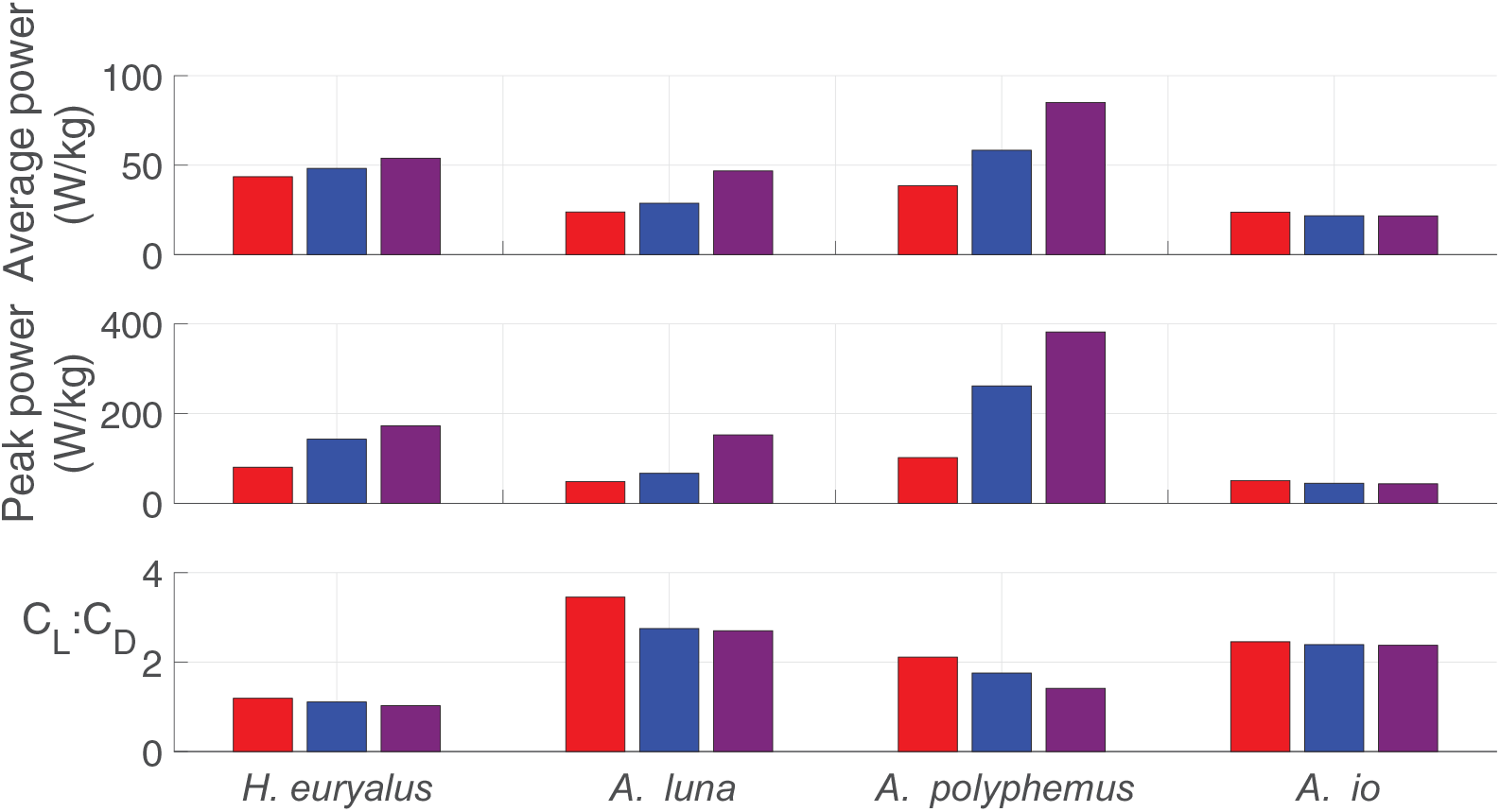
Bar plots showing the wingstroke-averaged and peak power dissipated in drag and the lift-to-drag ratio of the four silkmoth species *H. euryalus, A. luna, A. polyphemus* and *A. io*. Three colors of bar graphs correspond to the Models 1 (red), 2 (blue) and 3 (purple) of the body kinematic configurations.

## 4. Discussion

### 4.1 Body oscillations are associated with wingbeat frequency and wing loading

As established in some butterfly studies [6], [7], body oscillations are the consequence of the coupling between wing flapping and body dynamics. Because of the wing-body coupling dynamics, all flapping insects experience a degree of body oscillations driven at wingbeat frequencies. However, in butterflies and silkmoths, these oscillations are large and pronounced. Amplitudes of 25-27^°^ and mean values of 70-80^°^ are typical of forward flight (*<* 1 ms^−1^ flight speed) in butterflies *Kallima inachus* [21]. In this study, we observed amplitudes of up to about 40^°^ with a mean value of about 20^°^ in silkmoths *A. polyphemus* flying forward at 2-3 ms^−1^ with up to 1 ms^−1^ variation in vertical speed per wingstroke.

Butterflies and silkmoths have low wingbeat frequencies but large wings and flapping amplitudes to generate sufficient force for flight, compared to insects of similar body size and mass. Because the oscillation amplitude increases with the flapping time period and varies inversely with wing loading, the size of body oscillations is expected to be noticeably large in butterflies and silkmoths. Wing flapping excites the pitching dynamics of the body due to a time-varying pitching moment generated by both aerodynamic and inertial forces of wing flapping. A flapping flier with larger wings generates larger peak forces and thus a larger peak pitching moment. A flier with a smaller body mass has a smaller body moment of inertia and thus its body is prone to experiencing larger pitch accelerations. These larger peak accelerations, when applied periodically through lower wingbeat frequencies (and thus longer time periods), make the body pitch angle oscillate at larger amplitudes. As we show in our results, the product of wingbeat frequency and wing loading of silkmoths is negatively correlated with the amplitude of body pitch oscillations. Thus, silkmoths with slower and larger wings and smaller body mass show a larger magnitude of body oscillations. Even though the body pitch oscillations can be explained using a passive mechanism, the role of active abdominal flexion in partially inducing and controlling body pitch oscillations cannot be ignored due to the evidence for active abdominal flexion in moths and butterflies\ [7], [22], [23].

### 4.2 Wing inertial force alone cannot explain body oscillations

While silkmoths and butterflies are likely convergent in their body oscillations because the bombycoid moths ancestral to silkmoths likely had wingshapes more like hawkmoths (Sphingidae) wings. However, while both utilize oscillations, the mechanism for producing them seems different. In silkmoths, we found that both aerodynamic and inertial forces (Fig. 3) roughly equally contribute to the body oscillations in pitch and vertical velocity as opposed to birds and butterflies where wing inertial forces and aerodynamic forces, respectively, drive the body oscillations [6], [9]. Silkmoth wings are thin membranes covered in tiny scales made of chitin. Unlike birds, they lack bones or muscles within the wing itself and thus their wings are much less dense than bird wings. This implies that for wings of equal area and similar movement, a silkmoth wing would impart smaller inertial force but equal aerodynamic force on the body as compared to a bird wing. Thus, the wing inertial force in silkmoths contributes comparably to the effects of the aerodynamic force on the within-wingstroke dynamics of flight.

### 4.3 Body oscillations reduce power requirements of flight by reducing drag without compromising weight support

In steady forward flight, silkmoths (in this study) and butterflies pitch nose down during the down-stroke and nose up during the upstroke [23], [24]. This results in a time-varying stroke plane that remains steep during the downstroke and shallow during the upstroke [23]. This lets the fliers redirect the aerodynamic force into usable directions [24]. Precisely, a shallower stroke plane supports weight during the downstroke and a steeper stroke plane generates forward thrust during the upstroke by changing the direction of jet flow [21], [23]. Moreover, they can also control the direction of flight by controlling the direction of this jet flow. Thus, they can transition between forward and vertical flight by modulating the mean and amplitude of the body pitch oscillations [21]. However, it was not well known how the body oscillations affect the inclination angle of the wing and thus redirect the aerodynamic force into usable directions.

Compared to other insects of similar body size and mass, butterflies and silkmoths save power by using low wingbeat frequency. However, flapping larger wings with some amount of power being spent on exciting body oscillations raises the power requirements for flight. For both hover and forward-climbing trajectories of Monarch butterflies, the passive body oscillation arising from wing-body coupling results in a reduction of energy and power consumption [7]. In our study, we found that body oscillations in silkmoths, particularly oscillations in *χ*, can help maintain a low angle of attack *α*_v_ of the wing. This is achieved by body pitch oscillations operating at an appropriate phase relative to the flapping wing motion so that the incoming airflow is incident at a smaller angle on the wing thus maintaining a smaller *α*_v_, particularly during the downstroke. This results in smaller drag and thus reduces the aerodynamic power requirements of flight.

Even though reducing the power requirement of flight is beneficial, reducing drag in fliers like silkmoths that use larger stroke-plane angles during the downstroke and smaller stroke-plane angles during the upstroke can potentially reduce weight support and forward thrust due to a reduced overall aerodynamic force. However, body pitch oscillations can compensate for this effect by keeping the wings inclined at appropriate angles during each half-stroke. We found using our blade element model that how the wings turn due to the oscillations in *χ* can redirect the aerodynamic force vector during the downstroke and the upstroke to enhance weight support and forward thrust respectively. In the first half during the downstroke, a shallower body angle enforces a smaller *α*_h_ and hence, directs the aerodynamic force vector more vertically upward. Thus, most of the aerodynamic force during downstroke provides weight support rather than applying backward thrust on the body. For our species-averaged kinematics of silkmoths *A. luna* and *A. polyphemus, α*_h_ dips further and even stays negative for most of the downstroke duration. This makes the horizontal components of the aerodynamic force to be directed forward hence providing forward thrust even during the downstroke. In the latter half of the upstroke, a larger *α*_h_ of more than 50^°^ directs the aerodynamic force more horizontally forward which means most of the aerodynamic force generated during this half-stroke is utilized in propelling the body forward rather than pushing it downwards. This also enhances forward thrust and the overall weight support over the entire wingstroke. Body oscillations have likely independently evolved in both silkmoths and butterflies and have convergent aerodynamic consequences. Butterflies have also been shown to generate most of their weight support during the downstroke and propel themselves forward during the upstroke [21]. However, in the current study, we found that considerable forward propulsion can also be generated during the downstroke if the body tilts further nose-down to make the wing surface go below the absolute horizontal plane.

Besides the role of wing-body coupling, active abdominal flexions can also contribute to body pitch oscillations. A study on monarch butterflies has found that oscillatory abdominal flexions, which are more pronounced in butterflies, increase the magnitude of body oscillations [7]. This results in a reduction in mean power requirements of flight by up to 6%, providing further evidence for reduced power requirements due to body oscillations [7]. Active abdominal flexion in a hawkmoth and butterflies has been shown to have an active role in flight stabilization [22], [23], [25], [26]. Active articulation of the thoracic-abdominal joint redirects the aerodynamic force vector for effective flight control [22]. This force redirection is similar to what we refer to as force redirection arising from the wing rotation due to body pitch oscillations. Even though we have not explored the effect of this force redirection on flight stability, the body oscillations likely reduce the peak pitching moment over each half-stroke and hence reduce the total pitch impulse the body experiences in each half. This may provide a passive pitch stability mechanism in silkmoths during steady forward flight. Most passive stability analyses in the past have focused on high wingbeat frequency fliers that do not display passive body oscillations. Thus, our work also lays the foundation for this future study in silkmoths to directly test the role of body pitch oscillations in pitch stability.

## 5 Conclusion

Many silkmoths with low wingbeat frequencies and larger wings have an erratic flight behavior. Their bodies bob and pitch in a periodic fashion during steady forward flight. These body oscillations are coupled to their flapping wing motion and have strong aerodynamic implications. To explore which forces cause body oscillations and what are their aerodynamic consequences on flight performance, we recorded 3D forward flight kinematics of four silkmoth species flying forward in a wind tunnel. Then after extracting kinematic measurements and performing species-averaged aerodynamic fits of the four silkmoth species, we calculated the aerodynamic forces and power using a quasi-steady blade-element method to analyze the dynamics and performance of their flight. Given the observed patterns of kinematics, we found that their inertial and aerodynamic forces are equally dominant in generating body oscillations in linear velocity and body pitch angle. The oscillations in body pitch angle are associated with wingbeat frequency and wing loading of silkmoths and are coupled with the movement of flapping wings. This coupling exists in the form of a small range of phase difference between body pitch angle and wing sweep angle where flight power requirements are reduced by lowering the drag force on the wings. The oscillatory body pitch relative to periodic wing flapping is at a phase that maintains flight at a smaller angle of attack and redirects the aerodynamic force for enhancing upward and forward force. This reduces the total aerodynamic force by reducing the drag force without compromising weight support and forward thrust. Body oscillations might be ecologically important for silkmoths in two ways. They manifest as an erratic flight behavior and hence can help evade predators. Moreover, they reduce the aerodynamic power requirements of flight which is helpful because silkmoths rely on limited energy supplies as they do not feed as adults.

## 6 Acknowledgments

We would like to thank Hajime Minoguchi, Burhanuddin Bhinderwala and Cadence Brown for their help in data collection and digitization. We would also like to thank Megan Matthews, Varun Sharma, Joy Putney, Leo Wood, Ellen Liu and Ethan Wold for helpful discussions and feedback.

